# Micrometer-resolution reconstruction and analysis of whole mouse brain vasculature by synchrotron-based phase-contrast tomographic microscopy

**DOI:** 10.1101/2021.03.16.435616

**Authors:** Arttu Miettinen, Antonio G. Zippo, Alessandra Patera, Anne Bonnin, Sarah H. Shahmoradian, Gabriele E. M. Biella, Marco Stampanoni

**Affiliations:** Institute for Biomedical Engineering, University and ETH Zurich, Zurich, Switzerland; Swiss Light Source, Paul Scherrer Institute, Villigen, Switzerland; Institute of Neuroscience, Consiglio Nazionale delle Ricerche, Milan, Italy; Center for Cellular Imaging and NanoAnalytics (C-CINA), Biozentrum, University of Basel, Switzerland

**Author notes:** Present address of the author is Department of Physics, University of Jyvaskyla, Jyvaskyla, Finland (Arttu Miettinen). Present address of the author is National Institute of Nuclear Physics INFN, Turin, Italy. Corresponding authors: Arttu Miettinen, Antonio G. Zippo. Both corresponding authors contributed equally to this work.

## Abstract

Nervous tissue metabolism is mainly supported by the dense thread of blood vessels which mainly provides fast supplies of oxygen and glucose. Recently, the supplying role of the brain vascular system has been examined in major neurological conditions such as the Alzheimer’s and Parkinson’s diseases. However, to date, fast and reliable methods for the fine level microstructural extraction of whole brain vascular systems are still unavailable. We present a methodological framework suitable for reconstruction of the whole mouse brain cerebral microvasculature by X-ray tomography with the unprecedented pixel size of 0.65 μm. Our measurements suggest that the resolving power of the technique is better than in many previous studies, and therefore it allows for a refinement of current measurements of blood vessel properties. Relevant insights emerged from analyses characterizing the regional morphology and topology of blood vessels. Specifically, vascular diameter and density appeared non-homogeneously distributed among the brain regions suggesting preferential sites for high-demanding metabolic requirements. Also, topological features such as the vessel branching points were non-uniformly distributed among the brain districts indicating that specific architectural schemes are required to serve the distinct functional specialization of the nervous tissue. In conclusion, here we propose a combination of experimental and computational method for efficient and fast investigations of the vascular system of entire organs with submicrometric precision.

## 1. Introduction

Cerebral blood vessels sustain neuronal activity by providing metabolic components and oxygen to the brain tissues and by removing catabolic waste products. Specifically, it has been estimated that in humans and primates, synaptic activity and action potentials account for about 96% of the total energy consumed^1^, events enabled by a tight coupling among neuronal, glial and vascular cells. While much efforts have been spent to reconstruct in detail the sophisticated networks of specific neuronal circuits, no comparable achievements have solved the intricate microvascular architecture even in small nervous systems. Precise depiction of the vascular structure is also important for the comprehension of the biophysical mechanisms governing the crucial interplay among neurons, glial cells and blood vessels, a physiological event known as neuro-vascular coupling^2^.

The brain vascular structure is altered in many neurological diseases such as cerebrovascular diseases and various forms of Alzheimer’s and Parkinson’s diseases which constitute major health issues worldwide^3^. Many of these conditions are related to changes in the structure of the cerebral blood vessels, such as increased or decreased capillary density, microbleeds, stiffening of artery walls, or increase in vessel tortuosity^4,5^.

Reconstructing the structure of the whole vascular network in a brain is challenging as the whole brain is generally very large compared to the smallest microvessels. Two imaging techniques often applied to this purpose, in the context of mouse brain analyses, are magnetic resonance imaging^6^ and light sheet microscopy^7–9^. In these techniques resolution and pixel size are however often closer to 10 μm rather than 1 μm, and usually too large to unveil the smallest microvessels that have diameters down to less than 5 μm. On the other hand, X-ray tomographic microscopy (CT) offers higher resolution and the advantage of not requiring extensive sample preparation. Preparations such as tissue clearing increase the possibility of deformations in the sample, and their efficacy could vary spatially^10,11^.

Previously the cost of high resolution in CT was small sample size^12^. However, recent advances in imaging speed^13,14^ and post-processing algorithms^15^ allow for imaging of centimeter-scale samples in reasonable time (hours) with micrometer-scale resolution^16^. Notwithstanding, in all high-resolution imaging modalities capable of imaging large samples, the amount of image data to be analyzed and interpreted is very large, often in the range of terabytes. The large data size highlights the need to choose analysis algorithms and techniques such that the image analysis process can be performed in a reasonable time. Particularly favorable are algorithms that can be parallelized and distributed across computer clusters. Recently there have been several proposals of analysis software that can be applied to brain analysis^7,17,18^. Most of these are not openly available to the whole scientific community, are in early stage of development, or are geared towards single technique or imaging modality.

In this work, we show that synchrotron-based phase-contrast tomographic microscopy can be advantageously used to image micro-vessels in the whole mouse brain. In our technique the microvessels were perfused with contrast agent (Indian Ink^19^) and fixed with formalin solution. The perfused sample was tomographically investigated in a mosaic imaging mode, and the individual tiles were combined into one large volume image of the whole brain using a non-rigid stitching algorithm^15^. A main advantage of the mosaic imaging mode is that slow deformations of the sample do not lead to imaging artifacts, in contrast with more traditional CT imaging techniques. Finally, the full 11 TB volume image was segmented and analyzed using an in-house developed and freely available software, capable of processing the images in a few days using a small computer cluster as detailed below in the Section 4.

From the volume images we generated a vascular graph consisting of vessel branches and bifurcation points. The graph embedded various information related to the topology and morphology of the microvessel branches, e.g. length and average diameter. Subsequently, we determined the anatomical regions corresponding to our blood vessel space by registration of the volume image to the Allen mouse brain atlas^20^. Eventually, we described the vascular anatomy of the brain in different anatomical regions using quantities such as microvessel tortuosity, bifurcation density, vessel length density, and intercapillary distance.

## 2. Results

### 2.1. Sample preparation and imaging

Whole-brain samples were prepared by intracardiac perfusion of the vascular system with Indian Ink, according to the protocol in ^19,21^. The brain was then extracted and stored in polyphosphate buffered solution to maintain constant hydration until and during the data acquisition. The brain sample was CT imaged in a mosaic phase-contrast imaging mode where partially overlapping individual tomograms are taken side-by side to cover the whole volume of the brain. The tomograms were stitched into one large volume image using a non-rigid stitching algorithm^15^ that accounts for small deformations of the overlapping regions. The dimensions of the final volume image were approximately 13600 × 13600 × 30000 pixels with a pixel size of 0.65 µm. Finally, the volume image was denoised and segmented using standard image analysis algorithms.

### 2.2. Image quality

Initially, the quality of the volume image and the segmentation was visually evaluated by several imaging specialists. The algorithmic segmentation was found to visually match most of vessels identifiable on the volume image, see Figure 1 and Supplementary Animations 1-3. In order to find a quantitative estimate of the quality of the segmentation, an operator compared 2000 blocks (30×30×30 pixels each, from random locations) of the segmented image to the original. For each block, the operator determined whether there was a vessel in the block and whether it was segmented correctly. Confusion table (Supplementary Table 1) was calculated from operator’s answers. The segmentation agreed with operator’s perception of vesselness with sensitivity, specificity, F_1_-score and Youden’s J statistics of 0.943, 0.996, 0.960, and 0.939, respectively. Figure 1A-D shows various visualizations of the segmented microvessels.

**Figure 1.**
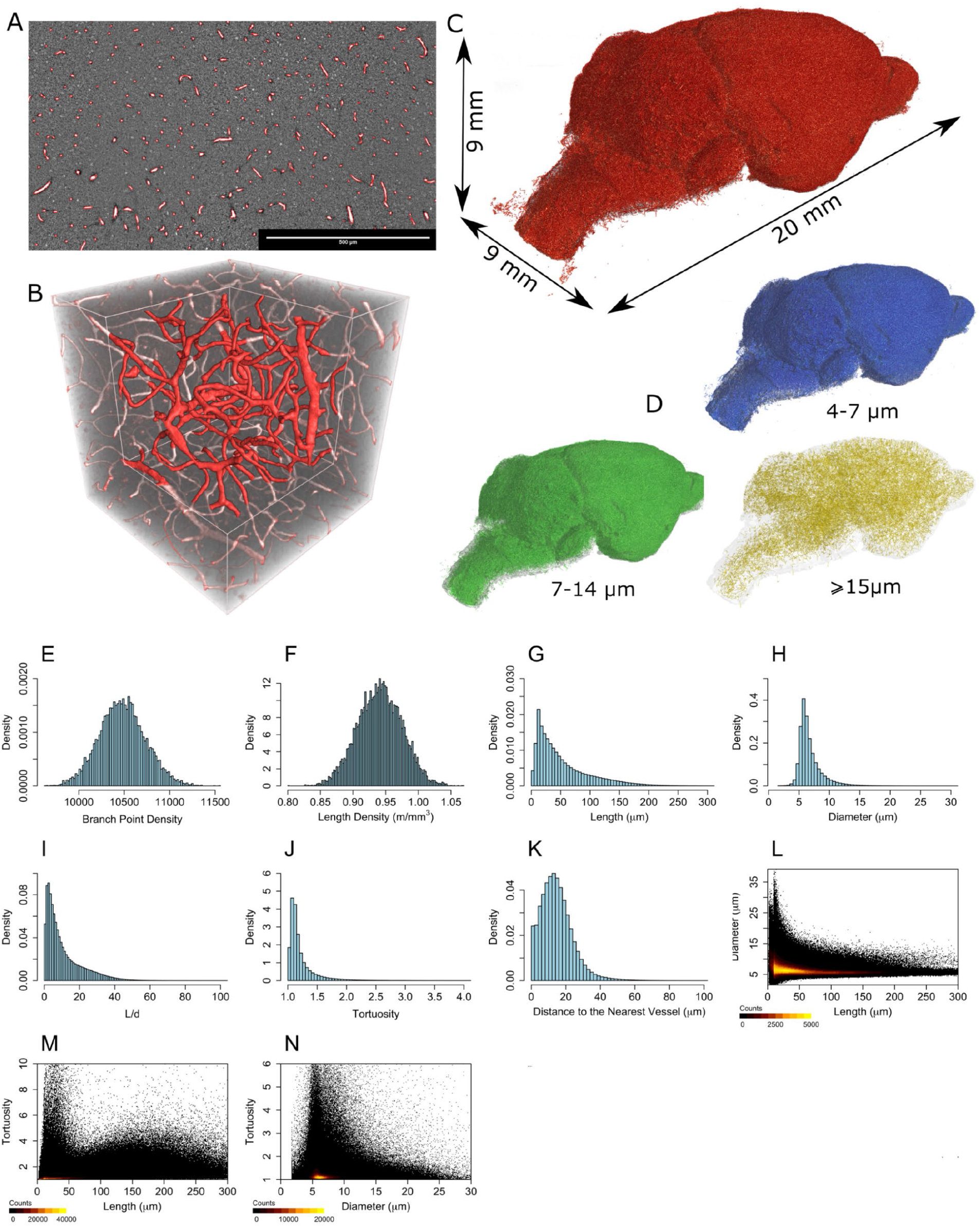
Visualizations of the segmented vessels (A-D) and uni- and bivariate distributions of various quantities measured from the segmented vessel data (E-N). A) Edges of segmented regions drawn as red outlines on top of a small part of the original non-segmented volume image. B) Visualization of the segmented vessels (red shade) overlaid on top of semi-transparent non-segmented data (grayscale). A volume of 325 μm × 325 μm × 325 μm is shown, and the corner nearest to the viewer has been cut away from the non-segmented data. C) Visualization of blood vessels in the whole tomographic image. D) Visualizations of blood vessels in specific diameter ranges.

Based on visual inspection of the blocks that were identified as badly segmented, the largest segmentation errors were found near large arteries and veins (diameters typically in the range of tens or hundreds of micrometers) that are not thoroughly perfused with the contrast agent (Indian Ink). We have not considered these vessels in the analysis. However, as shown by the statistics calculated from the confusion table, the number of such vessels is small compared to the microvessels, and therefore they do not contribute significantly to the results.

### 2.3 Overall structure of microvessels

In order to quantify the structure of individual microvessel branches, we found the centerline of each branch, and the bifurcation points of two or more branches. Additionally, we measured the length and the diameter of each branch.

We estimated the whole brain to contain approximately 4.3 million bifurcation points and 5.0 million vessel branches. Average bifurcation and length densities are 11000 ± 2000 per mm^3^ and 1.0 ± 0.1 m/mm^3^, respectively (Figure 1E-F), where the reported error limits correspond to uncertainty caused by the image analysis processes. The estimated length densities are somewhat higher than most previously reported values that are in range [0.440, 0.922] m/mm^3^ ^4,7,9,22,23^. Previous data on bifurcation density reported^7^ a value of approximately 3500 per mm^3^. The larger densities encountered in this study suggest that the true resolving power of the analysis pipeline used here is higher than in most of the previous studies. Additionally, different decisions made while choosing whether multiple nearby bifurcation points represent single physical bifurcation might lead to varying estimates of bifurcation density. Length density did not suffer from such an ambiguity in its definition and its value agrees better with the previously reported ones.

For the first time, our results returned estimations of the whole-brain vascular length (295 m), and of the average vessel branch length (53 ± 3 μm) (Figure 1G) at the level of the single microvessels. Average vessel diameter was 5.8 ± 0.4 μm in the whole brain (Figure 1H). It is conveniently between previously reported values of 4.25 μm^22^ and 8 μm^7^. The differences might be caused by various sample preparation routines such as perfusion and optical clearing. It was not certain that possible shrinkage or swelling of the vessels caused by these operations can be easily accounted for^24^, particularly when the focus is on local micro-scale properties and not on overall average deformation. In particular, any inaccuracy caused by local non-isotropic deformations are easily propagated to the results due to the small diameter of the vessels (in pixels, 5.8 μm = 8.9 pixels). According to results in Figure 1L vessel diameter was almost constant in the longer branches but varied more in the shorter ones.

The length over diameter ratio (L/d, dimensionless length) is related to the pressure loss in the vessel through the Darcy–Weisbach equation. Large values of L/d indicate that the vessel is long and thin, and the flow resistance of the vessel is large. We found an average L/d value of 11 ± 1 (Figure 1I) and largest values ranging to more than 100. The values indicate a large spectrum of putative flow resistance regimes^25^.

Tortuosity is a measure of how much a blood vessel segment twists, with high values typically related to pathologies^26^. Average tortuosity of the vessels was 1.24 ± 0.01 (Figure 1J), a value which was maximized with 20-30 µm length vessels and generally increases with the vessel length (Figure 1M). Further, tortuosity was highest in vessels of diameters between 5 to 7 µm (Figure 1N), indicating that the purpose of the vessels in this size range is to assist in even transport of metabolic components and waste products to and from the tissue, in contrast to efficiently transferring them for long distances. The average distance the products must transport outside of blood vessels equals to the distance to the nearest microvessel, and that was measured to be 15 ± 1 μm. The value corresponds to approximately 2.5 average microvessel diameters (Figure 1K).

### 2.4 Differences in the vascular structure between anatomical regions

In order to obtain a more detailed picture of the structure of the vessel network, we co-registered the obtained mouse brain vessels with the Allen mouse brain atlas at the highest resolution of the atlas (100 μm pixel size). In addition, we clustered the atlas brain regions into two different hierarchical groups such that very small regions were combined in order to guarantee that each clustered region contained statistically significant number of vessel branches. In the first (finer) level, we have 44 different regions (Supplementary table 1, left column, *regions*), and in the second coarser level 11 regions (Supplementary table 1, right column, *macroregions*).

We calculated the vessel measures in each region (Figure 2, Supplementary figure 1, Supplementary animations 4-11), and non-parametric Kruskal-Wallis test indicated that results for all the regions do not come from the same distribution (*P*<machine precision). This was true for all the measures. In particular, the branch point density was statistically different among the macroregions (Figure 2B) and regions (Supplementary figure 1B) as well as the length density (Figures 2C, Supplementary figure 1C). The vessel length was non-uniformly distributed among macroregions (Figure 2D) and regions (Supplementary figure 1D). Approximately 12% of all the vessel branches connected two or more anatomical regions, and the rest remained inside single region.

**Figure 2.**
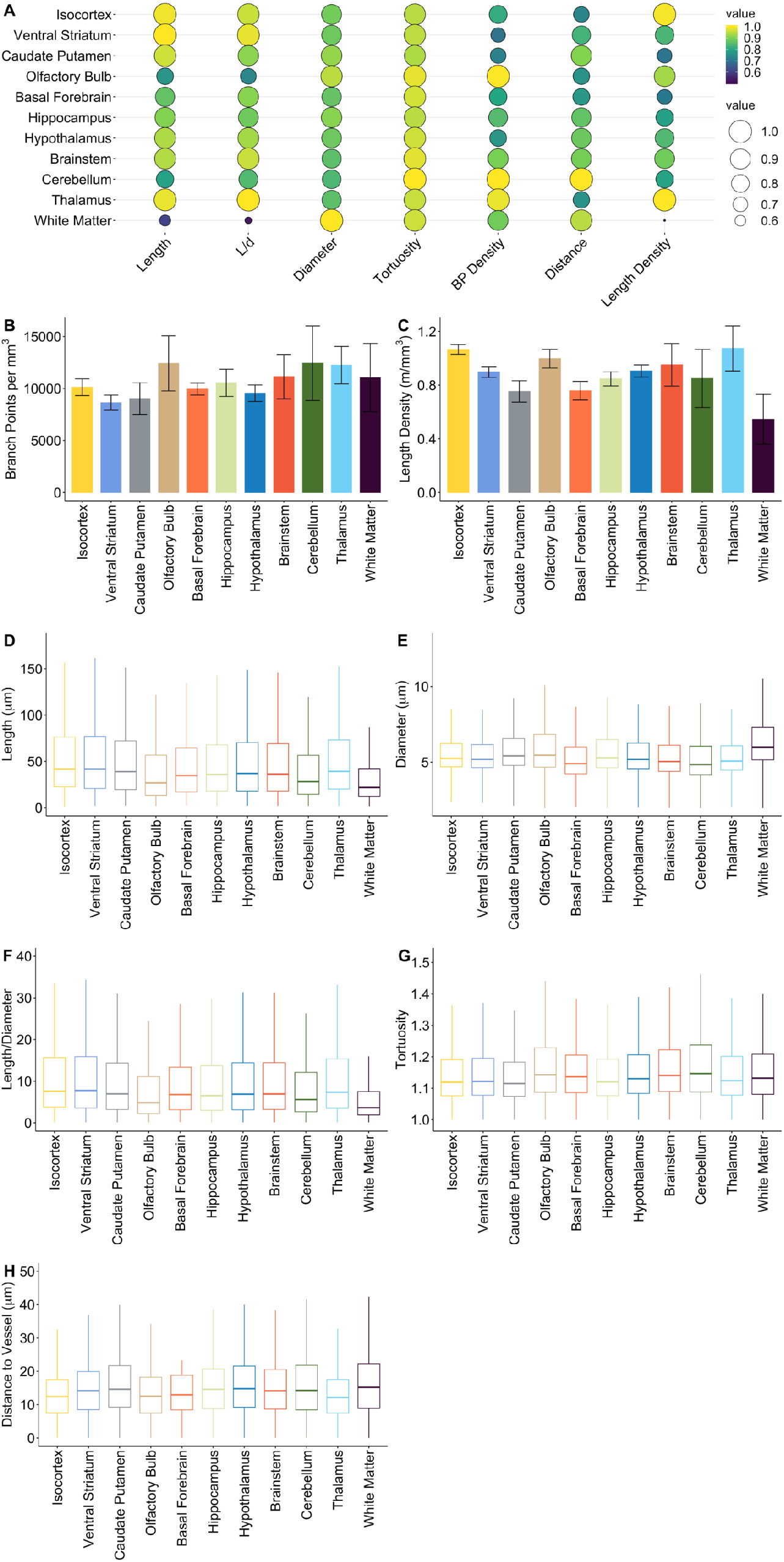
A) Bubble plot showing correlations between various measured quantities in different anatomical macroregions. B-H) Values of various measured quantities in different anatomical macroregions. In B and C the error limits describe uncertainty caused by the image analysis process.

Post-hoc significant differences of the various measured quantities (Tukey tests, Supplementary files) composed a hierarchical descending order (*Hasse diagrams*^27^, Figure 3) for the macroregions, where statistically significantly different regions are placed on different levels of the hierarchy. These orderings highlighted the Ventral Striatum and the Isocortex as the macroregions with longest vessel branches and, oppositely, the White Matter and the Olfactory Bulb as the macroregions with shortest vessel branches. In terms of vessel diameters (Figures 2F and 3), the White Matter and the Olfactory Bulb were the macroregions with largest values and the Cerebellum, while the Brainstem and the Thalamus had the smallest vessel diameters. The L/d ratio pointed (Figures 2G and 3) the Hypothalamus and the Ventral Striatum as the macroregions with the highest values, the White Matter and the Olfactory Bulb were instead the macroregions with the smallest L/d ratio. The tortuosity reached the highest values (Figures 2H and 3) in the Thalamus and the Ventral Striatum while the smallest values were estimated in the White Matter and the Olfactory Bulb. The distance to the nearest vessel was highest (Figures 2I and 3) in the Cerebellum, the Brainstem, the Caudate Putamen and the White Matter. Conversely the distance reached the minimum values in the Isocortex, the Olfactory Bulb and the Thalamus.

**Figure 3.**
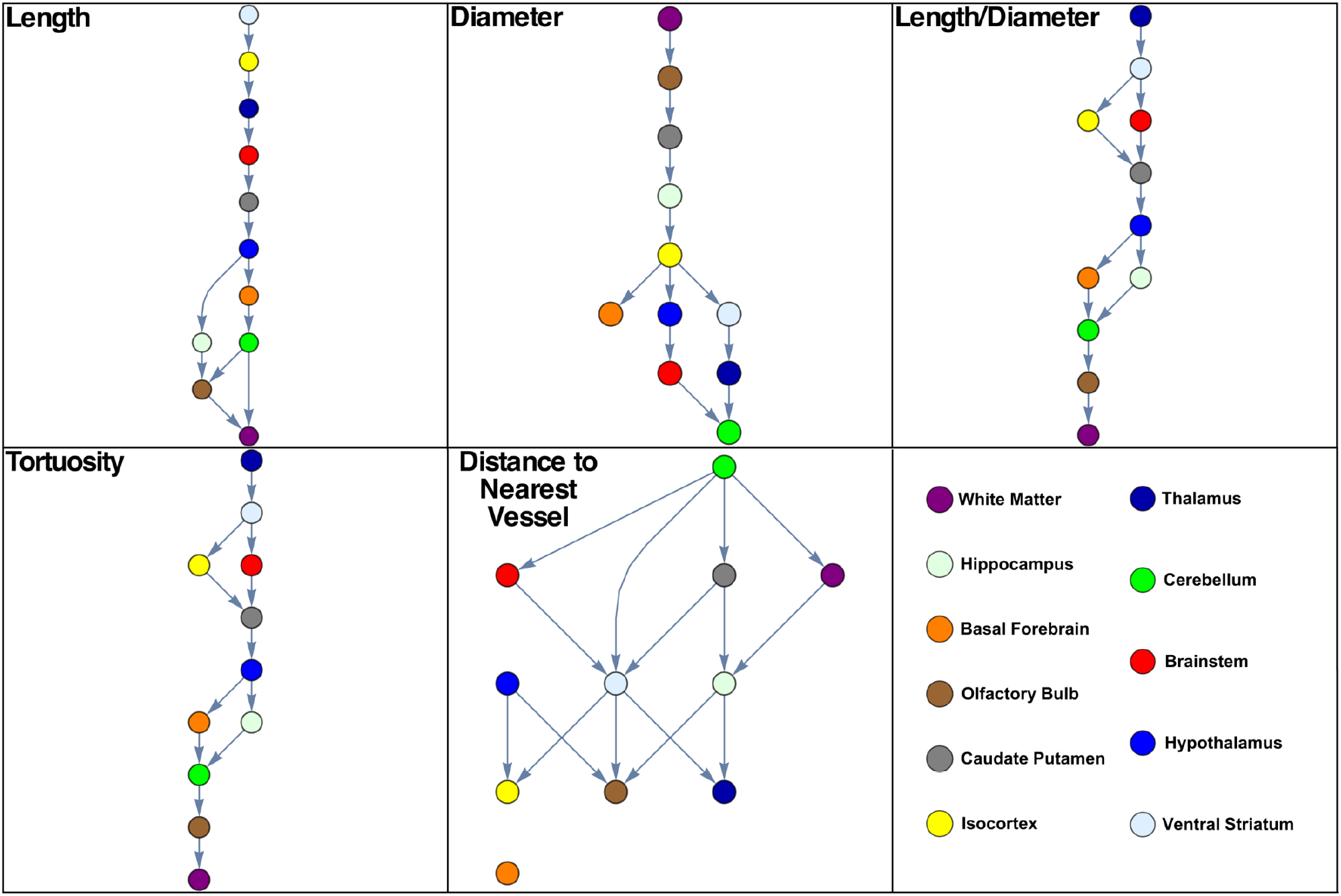
Hasse diagrams showing statistically significant differences in the different macroregions and different quantities. The regions are placed on different levels if the difference in the corresponding quantity is significant between the regions. The topmost levels correspond to the largest values of the quantities.

We did similar analyses for the 44 finer anatomical regions (Supplementary figure 1). Remarkably, vessel lengths were longest in the Thalamus subregions and in the Extrapyramidal Tract, and shortest in the Frontal Pole (Figure S14). Vessel diameter was largest in the Accessory Olfactory Bulb and in distinct White Matter regions (Optic Nerve, Corpus Callosum, Corticospinal Tract) while it was smallest in the Primary Somatosensory and Motor Cortices and the Retrosplenial Area (Figure S16). Eventually, the L/d ratio was greatest in the thalamic regions while the Frontal Pole, the Perirhinal Area the Olfactory Bulb, the Visceral and the Orbital Area were characterized by smaller values (Figure S17). In conclusion, geometrical, morphological and topological features of mouse brain microvessels appeared regionally specific suggesting distinct roles in support of local specialization of brain districts.

### 3. Discussion

The results highlighted peculiar characteristics of specific macroregions that were mainly the white matter, the olfactory bulb and the cerebellum but also the striatum and the somatosensory, motor and visual cortices, which appeared to get extreme values in our estimations. Indeed, we observed a strong correlation between the neuronal density^28,29^ and the numbers of branch points and tortuosity (Figures 4A and K), a weaker but sustained correlation has been detected also with the distance to the nearest vessel (Figure 4N). In addition, comparable and in some cases even stronger correlations held also in the glial cell density (Figure 4, third and fourth columns). Note that the estimations were biased by the zero neuronal density of white matter and, oppositely, by the high neuronal density of cerebellar layers (more than fivefold the average of other regions). Indeed, removing these two regions from the linear regressions results in stronger correlations between neuronal density and many of the measured quantities (Supplementary figure 2). Such postliminary considerations suggest that no trivial rules govern microvascular features in relation to the other existing cellular families (neurons and glia). Surprisingly, vessel tortuosity appeared to be the best predictor (as for linear regression) for neuronal density while the distance to the nearest vessels played the same role for glial density.

**Figure 4.**
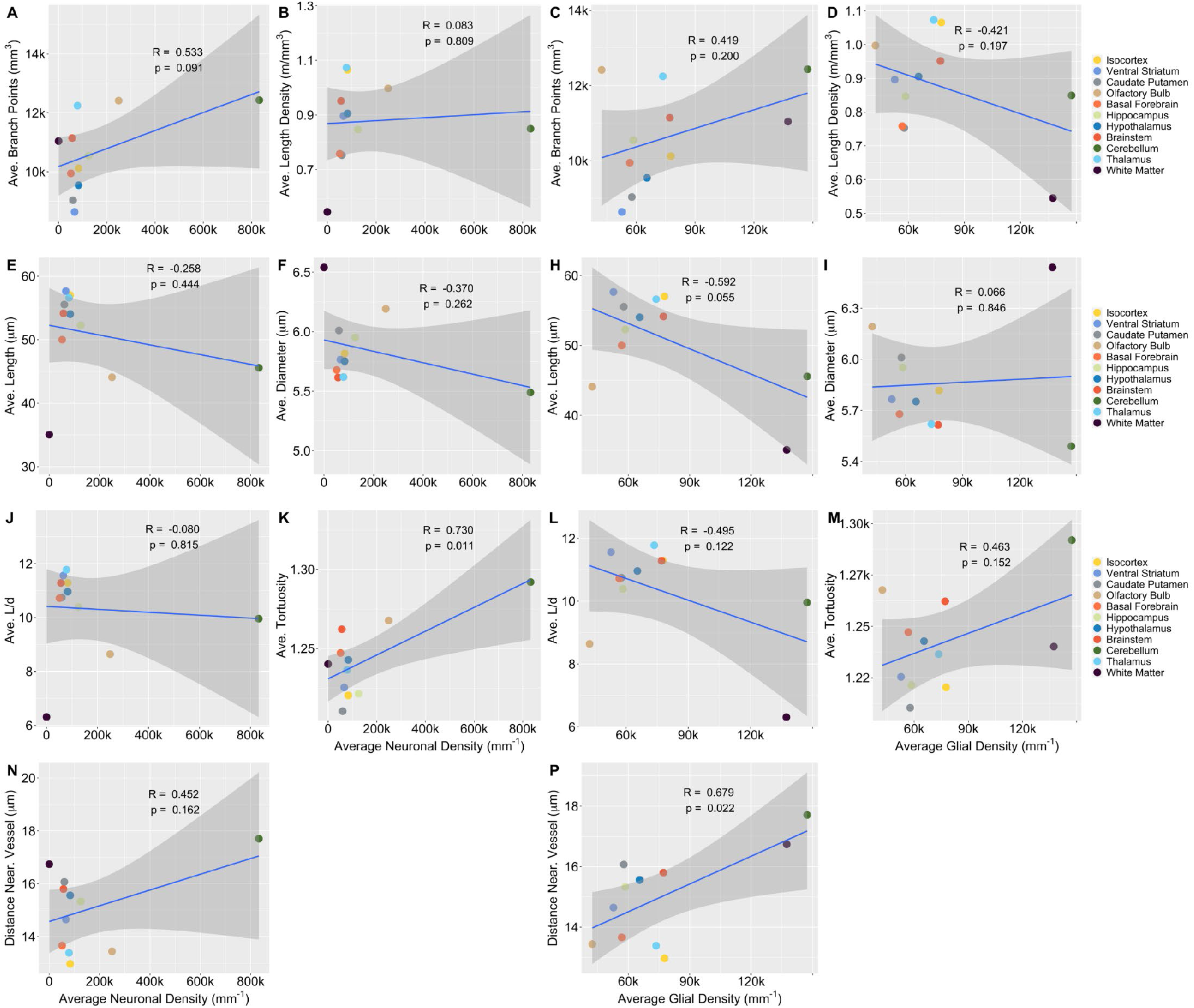
Correlations between average neuronal (first and second column) and glial (third and fourth column) densities and various measured quantities. Density data is from the EPFL mouse brain atlas^28^. Ave., average; Near., nearest.

Besides density correlations, a reasonable observation was that typical high energy demanding regions were denser of vessels with long segments and small diameters. A set of neurophysiological and anatomical considerations supported the observed results. The brain white matter is mostly composed by myelinated neuronal axons and glial cells. From metabolic perspective, axonal segments demand (with the complicity of astrocytes^30^) approximately 3-fold lower energies than their terminals^1,31^, albeit these estimations have been calculated on the amount of mitochondrial ATP consumption. Accordingly, although authors reported that oligodendrocytes, abundantly populating white matter^29^, provide metabolic support to neurons through monocarboxylate transporters^32,33^, our results showed low vascular density in the white matter regions mostly characterized by the presence of large vessels in terms of diameter.

The olfactory bulbs are important districts for the olfactory information processing, crucial for rodent behavior and survival. The olfactory bulbs are the second neuron densest brain structure^29^ (~250000 neurons per mm^3^). In olfactory bulbs tortuosity was very high (the second highest) indicating that vessels are highly twisted and curved. Vessels in this region were characterized by the highest branch point density, low segment length and large diameter.

The cerebellum is an evolutionary ancient three-layered brain section distinguished by the highest neuronal density of about 830000 neurons per mm^3^. As expected, our estimations showed that cerebellar vessels were the smallest in terms of diameter and one of the shortest according to segment length. Nonetheless, cerebellar vessels had the highest distance to nearest vessel among regions, an unexpected result which suggested that the ratio of vascular endothelial cells to the number of neurons is relatively low in comparison to other regions^28^.

The striatum is a subcortical region responsible for motor functions with a relatively low neuronal density^28,29^ (~64000 neurons per mm^3^), and is generally divided in its ventral and dorsal (caudate putamen) parts. Ventral striatum (~66000 neurons per mm^3^) had the longest vascular segments and relatively low diameters, compatible with a moderate level of energy demand.

At last, the neocortex, which is responsible of most of high-level brain functions and is characterized by a moderate neuronal density (approximately 83000 neurons per mm^3^), did not stand out in any vascular feature. This result seems to be in accordance with recent 18FDG-PET measurements of the homogeneous metabolism of the mouse cerebral cortex among other hindbrain and forebrain structures^34^.

As a summary, we presented a methodological framework for comprehensive and precise reconstruction of the entire microvasculature of the mouse brain at the unprecedented pixel size of 0.65 µm. Local synchrotron-based X-ray phase-contrast tomography combined with an attainable computational pipeline resulted in an effective methodology to investigate geometrical, morphological and topological features of vascular systems of *ex vivo* organs at their finest structure.

The approach proposed in this paper has a few limitations. First, the whole brain sample analyzed in this work was imaged in approximately 1200 tiles that required image acquisition session lasting more than 57 hours. Each tile was therefore imaged in approximately three minutes. The sample must be steadily mounted and stable such that during each three-minute interval it moves less than one pixel (0.65 µm), or otherwise the tomographic reconstructions of the individual tiles may contain artifacts. Although this requirement could still rise problems in various experimental setups, it is much easier to achieve than similar stability over the whole imaging session. Note that deformations between neighboring tiles are acceptable in the stitching method used here^15^. Second, the results shown here are based on a single animal and, although related literature does not indicate important variations in the brain microvascular architecture, conclusions of this work could be slightly different in a larger animal sample. Third, it seems to be hard to perfuse all vessels adequately with the proposed contrast agent (Indian Ink), and therefore the non-perfused vessel branches are missing from the analysis. This leads to biased results especially for the largest vessels, but according to results shown in Section 2 and Supplementary Table 1 the smaller vessels seem to be unaffected.

In the future, the methods proposed in this work can be used, e.g., to construct an atlas of microvessel geometry in mouse brain, both in healthy and pathological conditions, or to study blood flow in more detail using image-based flow simulations either in direct image-based modality^35^ or using the generated vascular graph^36^. In conclusion, we demonstrated that it is possible to make high-quality tomographic images of very large, fragile, moisture- and radiation sensitive samples, and analyze their structure with image-based measurements. We believe that the methodology introduced here generalizes well to many kinds of biological and engineered samples, and is particularly useful in cases where optical clearing required in many other imaging modalities is not possible or desirable.

## 4. Materials and Methods

### Mouse sample preparation

The experimental procedure was approved by the local veterinary authority of Canton Zürich, Switzerland (license number ZH184/2015).

After the loss of any reflex, before the death by the barbiturate overdose, the animals were prepared for the perfusion by the opening of the sternal plate and the thoracic cage. The beating heart was then gently clamped with flat tweezers and an 8-gauge metal needle with the smoothed tip was inserted into the left heart ventricle. The intracardiac perfusion was performed in a few stages: first with a Ca+/Mg+-free phosphate buffered saline (PBS) (100 ml, 37 °C), followed by a 4 % paraformaldehyde and Karnovsky’s fixative (100 ml, 37 °C), and then perfused with Indian Ink (50 ml, 40 °C), and finally with Karnovsky’s fixative (20 ml, 4 °C). A clear sign of the complete perfusion was the generalized blackening of all the mucosae, of nude skin surfaces (such as the snout, the paws) the thoracic viscera and the eyes.

Subsequently, the animals underwent euthanasia in order to extract the whole brain. The extracted brains were immersed in PBS solution and maintained in constant hydration conditions until and during the data acquisition.

### X-ray tomographic microscopy

Samples were imaged at the TOMCAT beamline of the Swiss Light Source at Paul Scherrer Institute (Switzerland). The dataset consists of 9 × 9 × 15 tomograms of 2048^3^ pixels each. The individual tomograms form an image mosaic, where overlap between neighboring images is 30% of their diameter in directions perpendicular to the rotation axis, and approximately 10% in the direction parallel to the rotation axis.

Each tomogram was reconstructed from 1001 X-ray projection images with the GridRec algorithm^37^. Paganin phase retrieval method^38^ was used before reconstruction. The projection images were acquired with 20 keV monochromatic X-ray beam, 0.65 μm pixel size, and 50 ms exposure time. The sample to detector distance was set to 100 mm. The total acquisition time was approximately 57 h.

### Stitching

We stitched the individual tomograms into one large volume image using a non-rigid stitching algorithm^15^. There, the locations and the orientations of the individual tomograms are globally optimized such that disagreements between them in the overlapping regions are minimized. Furthermore, the overlapping regions are deformed such that any remaining disagreements are eliminated. This processing ensures that the microvessels are continuous across boundaries of individual tomograms even in cases where the sample has deformed during image acquisition. The size of the stitched volume image was approximately 13600 × 13600 × 30000 pixels (11 TB with 16-bit pixels).

### Registration to the Allen atlas

The stitched volume image was downsampled to similar size than the annotated Allen adult brain atlas at its full resolution (version CCF 2017)^20^. The Atlas was then registered with the volume image, initially by an affine transformation, and further refined with a non-rigid B-spline transformation.

### Image segmentation

The stitched volume image was denoised using bilateral filtering^39,40^ (spatial σ = 1.3 μm, radiometric σ = 7.6% of full dynamic range), followed by high-pass filtering to remove large-scale intensity variations (spatial σ = 6.5 µm). The filtered image was segmented with a region-growing approach. To that end, initially all pixels whose value were above a threshold were classified as vessels. The vessel regions were grown until all bordering pixels had a value below a second threshold. All other pixels were classified as background. The threshold values were selected such that the segmentation visually corresponded to the vessels.

Possible gaps in the segmented vessels were eliminated by applying a morphological closing filter (radius = 3.25 μm). In addition to blood vessels, the segmentation process identified choroid plexuses, some small and separate non-vessel regions, and many planar structures (at the surfaces of the brain, mostly caused by remaining phase contrast artifacts) as vessels. In order to eliminate the small non-vessel regions, all foreground objects less than 685 μm^3^ (equivalent to 2500 pixels) in volume were discarded. As the blood vessels form a continuous network, this process does not have any effect on them. The choroid plexuses in the ventricular cavities were removed by masking the original segmentation with a mask where choroid plexuses were not visible. The mask was generated using morphological opening and closing operations.

Planar structures were eliminated by calculating a surface skeleton^41^ of the foreground, and eliminating all surfaces consisting of more than 5000 pixels. The surface skeleton was then refined into a line skeleton^41^ where each blood vessel is turned into a single pixel thick line located in the middle of the vessel.

### Image analysis

The line skeleton was traced in order to produce a graph representation of the microvessel network. In the graph, vessel bifurcation points are represented as vertices and vessels as edges. The bifurcation points were found and the center lines between them, representing individual branches of the blood vessel network, were traced in order to produce a graph representation of the microvessel network. For each bifurcation point, the corresponding anatomical region was recorded based on the annotated volume registered with the image (see Section “Registration to the Allen atlas”). For each microvessel branch, the length L of the branch^42^, distance D between its end points, and the cross-sectional area A of the vessel was recorded. The cross-sectional area was measured by taking two-dimensional cross-sectional slices of the vessel and measuring its area from those^43^. The effective diameter of the vessel was then determined as 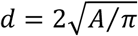, tortuosity as *L/D*, and slenderness as *L/d*. Branches shorter than 9.75 μm (equivalent to 15 pixels) and not connected to multiple other branches in both ends did not correspond to vessels and were pruned. Finally, the distance between the microvessels was quantified by calculating a distance map^44^ where each non-vessel pixel is associated the distance to the nearest blood vessel.

### Uncertainty analysis

The uncertainty limits were estimated using a Monte Carlo method, where the image analysis process is repeated several times with perturbed input parameters and the uncertainty limits are calculated from the distribution of the results. In order to speed up processing, uncertainty analysis was done on 58 blocks of the original volume image, 1500^3^ pixels each, selected randomly from all anatomical regions in the brain.

The values of the input parameters were drawn randomly from normal distributions with means given by the values used for analyzing the whole volume image, and standard deviations of 10% of the mean, except for radiometric σ and threshold values where 5% and 2.5% were used, respectively. The values of the standard deviations were chosen such that the segmentation result calculated with any single parameter perturbed by two standard deviations had visibly low quality. Total of 25 iterations were made for each block, and the average relative error for each output quantity was calculated. The relative error averaged over all the blocks (separately for each reported quantity) was applied in reporting the uncertainty limits for the full volume image.

### Statistics

Distributions of various quantities in different anatomical regions were calculated using simple statistical binning, or alternatively visualized using box plots showing the minimum, the maximum, the median, and the first and the third quartiles.

One-way comparisons of vessel measures among anatomical regions (or macroregions) were computed by using the non-parametric Kruskal-Wallis test. Subsequent pairwise comparisons were estimated with the Tukey post-hoc test. In all statistical tests in Figure 2 and Supplementary Figure 1, the Kruskal-Wallis test returned a p-value smaller the machine precision (2.16× 10^−16^).

### Software and data availability

Stitching, image segmentation and analysis was performed using an in-house developed software ‘pi2’, available at github.com/arttumiettinen/pi2. The software allows user-transparent distribution of the image analysis tasks on a computer cluster. We used a heterogeneous cluster with 10-30 available compute nodes, each equipped with 24-36 Intel Xeon cores and 180 GiB of random-access memory available to the analysis software. Depending on the availability of the resources, the analysis of the whole volume image takes approximately 5-10 days.

Registration with Allen atlas was done manually using the 3D Slicer software^45^ available at www.slicer.org, employing Transform, Landmark Registration, and Resample Image modules.

The visualizations and the supplementary animations were generated with MeVisLab, ImageJ^46^, Blender, Inkscape, and Gimp, available at www.mevislab.de, imagej.nih.gov/ij, www.blender.org, inkscape.org, and www.gimp.org, respectively. The statistical analysis was done in the R environment (www.r-project.org).

The image data and image analysis code is available at the PSI data repository^1^. The supplementary animations are available at YouTube^2^.

## Acknowledgements

This work was supported by Swiss National Science Foundation Grants CR23I2-135550 and 310030-153468. The authors acknowledge Bruno Weber and Ladina Hösli for providing laboratory resources for sample preparation.

## Author contributions

A.M. and A.G.Z. developed the image analysis methods, performed statistical analysis, created figures and supplementary visualizations and wrote the manuscript. A.P., S.H.S and A.B. developed sample preparation and imaging protocols and performed the CT experiments. M.S. and G.E.M.B supervised the project. All the authors participated in the finalization of the manuscript.

## Supplementary material

**Supplementary Table 1.** Confusion table for segmentation quality. The table was compiled by manually comparing the segmented image to the original in 2000 random locations.

|  |  | Actual class |  |  |
| --- | --- | --- | --- | --- |
|  |  | Vessel | Not vessel | Total |
| Predicted class | Vessel | 279 (14.0%) | 6 (0.3%) | 285 (14.3%) |
|  | Not vessel | 17 (0.9%) | 1698 (84.9%) | 1715 (85.8%) |
|  | Total | 296 (14.8%) | 1704 (85.2%) | 2000 (100%) |

**Supplementary Table 2.** Allen atlas regions corresponding to the coarse (macroregions) and fine level (regions) clustering.

| **Fine level Atlas region clustering** | **Coarse level Atlas region clustering** |
| --- | --- |
| Gustatory areas | Isocortex |
| Prelimbic area |  |
| Frontal pole |  |
| Orbital area |  |
| Visceral area |  |
| Perirhinal area |  |
| Agranular insular area |  |
| Infralimbic area |  |
| Temporal association area |  |
| Posterior parietal association area |  |
| Ectorhinal area |  |
| Anterior cingulate areas |  |
| Somatomotor areas |  |
| Somatosensory areas |  |
| Visual areas |  |
| Auditory areas |  |
| Retrosplenial area |  |
| Piriform area |  |
| Striatum | Ventral Striatum |
| Dorsal peduncular area |  |
| Pallidum |  |
| Cortical subplate | Caudate Putamen |
| Cortical amygdalar area |  |
| Piriform-amygdalar area |  |
| Postpiriform transition area |  |
| Accessory olfactory bulb | Olfactory Bulb |
| Anterior olfactory nucleus |  |
| Main olfactory bulb |  |
| Lateral olfactory tract nucleus | Basal Forebrain |
| Taenia tecta | Hippocampus |
| Hippocampal formation |  |
| Hypothalamus | Hypothalamus |
| Midbrain | Brainstem |
| Pons |  |
| Medulla |  |
| Cerebellum | Cerebellum |
| Dorsal thalamus | Thalamus |
| Ventral thalamus |  |
| Posterior thalamus |  |
| Anterior thalamus |  |
| Extrapyramidal tract | White Matter |
| Corpus callosum |  |
| Corticospinal tract |  |
| Optic nerve |  |

**Supplementary figure 1.**
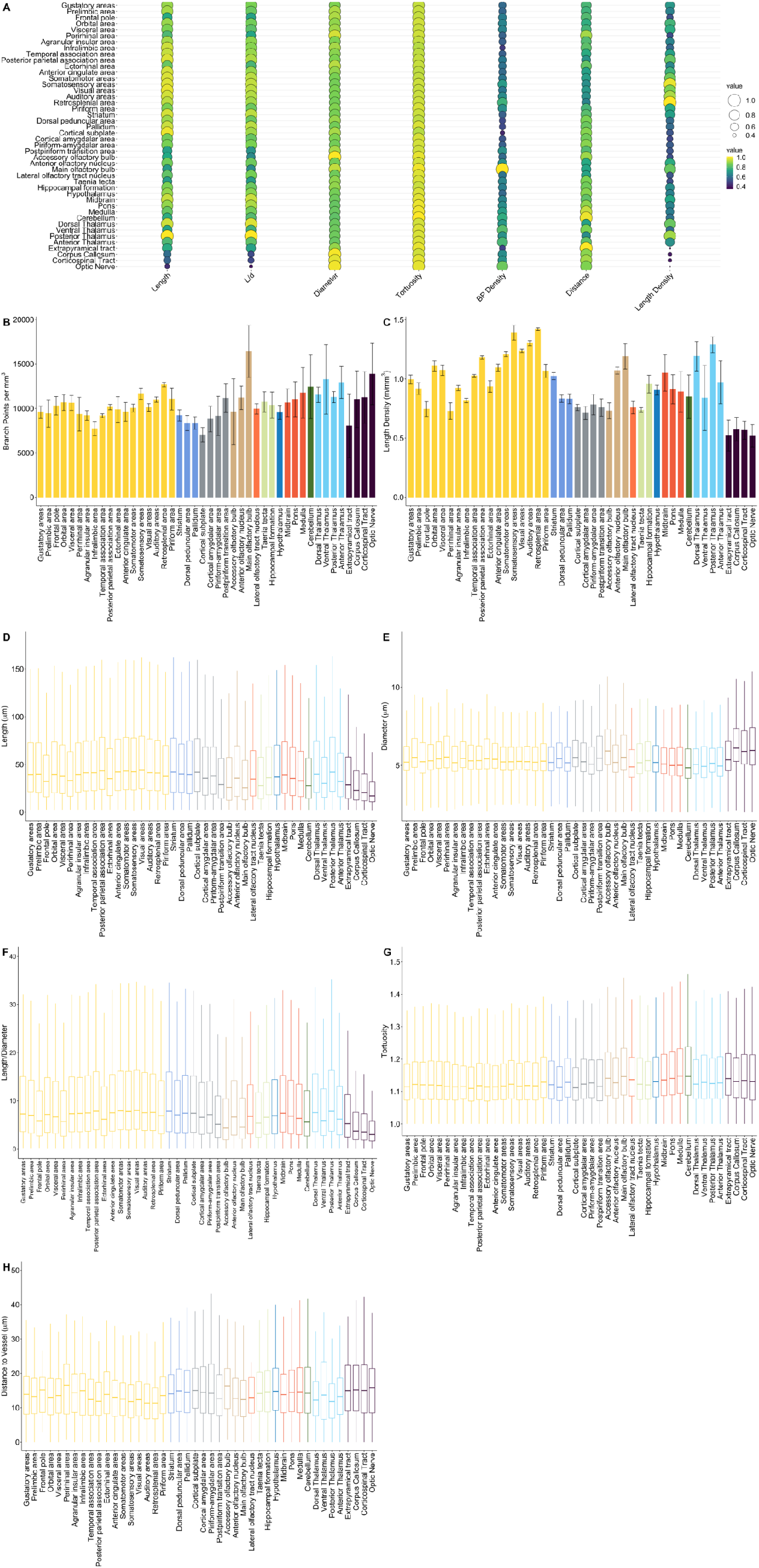
A) Bubble plot showing correlations between various measured quantities in different anatomical regions. B-H) Values of various measured quantities in different anatomical regions. In B and C the error limits describe uncertainty caused by the image analysis process.

**Supplementary figure 2.**
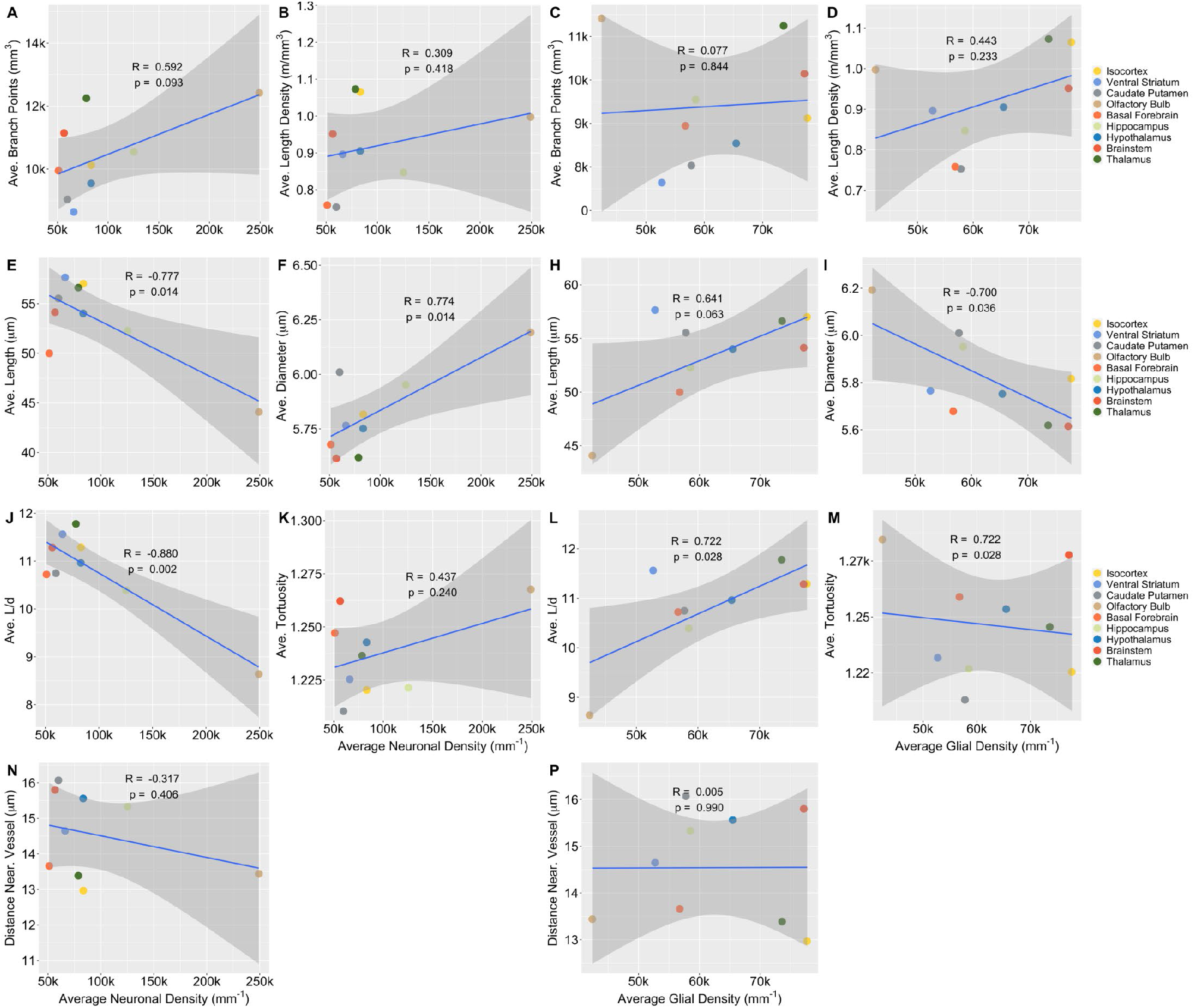
Correlation analyses between neural (first and second column) and glial (third and fourth column) densities and vessel measures as in Figure 4 but excluding the two outlier macroregions, the white matter and the cerebellum.

**Supplementary file 1**

Regional average differences and post-hoc statistics (Tukey test) of the vascular features for macroregional and regional comparisons.

**Supplementary animation 1**

Supplementary animation 1 is a visualization of the microvessels in the whole mouse brain as a maximum intensity projection of the original (non-segmented) volume image. The animation begins from a view where the whole brain is visible and zooms in until the individual microvessels are well visible. Available at https://www.youtube.com/watch?v=gbXkBWnqBs8.

**Supplementary animation 2**

Supplementary animation 2 visualizes the performance of the proposed segmentation pipeline. The edges of the segmented microvessels are drawn with red color over a slice through the original volume image. The animation begins from an arbitrary location inside the cerebrum and proceeds towards the cerebellum with a speed of 13 μm/s.

Available at https://www.youtube.com/watch?v=6mdv0gB1drE

**Supplementary animation 3**

Supplementary animation 3 shows a 3D visualization of the segmented vessel network.

Available at https://www.youtube.com/watch?v=2Znl3indW-8.

**Supplementary animations 4-11**

Supplementary animations 4-11 visualize spatial variations in the measured quantities between anatomical macroregions.

|  | Title | Available at |
| --- | --- | --- |
| 4 | Overview | https://www.youtube.com/watch?v=4hhurq0Fwx0 |
| 5 | Length density | https://www.youtube.com/watch?v=u7fjgwx2pbo |
| 6 | Bifurcation density | https://www.youtube.com/watch?v=sBXUp0d30WE |
| 7 | Branch length | https://www.youtube.com/watch?v=0eMp2ZGZc6M |
| 8 | Length/diameter ratio | https://www.youtube.com/watch?v=6cLs_9G8rXk |
| 9 | Distance to nearest vessel | https://www.youtube.com/watch?v=si4qJ-aO1sw |
| 10 | Vessel diameter | https://www.youtube.com/watch?v=NVKxlmkqzII |
| 11 | Tortuosity | https://www.youtube.com/watch?v=R92zNhzcSvw |

## Footnotes

1 https://doi.org/10.16907/1237d208-9057-4755-8049-40ee7a199b15

2 www.youtube.com/channel/UCPKtwMW6rfirNPHRNwNa_0A

